# The heparin-binding domain of VEGF165 directly binds to integrin αvβ3 and plays a critical role in signaling

**DOI:** 10.1101/2023.11.14.567104

**Authors:** Yoko K. Takada, Jessica Yu, Xiaojin Ye, Chun-Yi Wu, Brunie H. Felding, Masaaki Fujita, Yoshikazu Takada

**Author notes:** Corresponding Author: Yoshikazu Takada, MD, PhD, Department of Dermatology, Biochemistry and Molecular Medicine, UC Davis School of Medicine, Research III, Suite 3300, 4645 Second Avenue, Sacramento, CA 95817. <u>Masaaki Fujita</u>: Division of Clinical Immunology and Rheumatology, Kansai Electric Power Hospital, 2-1-7 Fukushima, Fukushima-ku, Osaka 553-0003 Japan.

## Abstract

VEGF-A is a key cytokine in tumor angiogenesis and a major therapeutic target for cancer. VEGF165 is the predominant isoform and is the most potent angiogenesis stimulant. VEGFR2/KDR domains 2 and 3 (D2D3) bind to the N-terminal domain (NTD, residues 1-110) of VEGF165. Since removal of the heparin-binding domain (HBD, residues 111-165) markedly reduced the mitogenic activity of VEGF165, it has been proposed that the HBD plays a critical role in the mitogenicity of VEGF165. Integrin αvβ3 has been shown to bind to VEGF165, but the role of integrin αvβ3 in VEGF165 signaling are unclear. Here we describe that αvβ3 specifically bound to the isolated HBD, but not to the NTD. We identified several critical amino acid residues in HBD for integrin binding (Arg-123, Arg-124, Lys-125, Lys-140, Arg-145, and Arg-149) by docking simulation and mutagenesis, and generated full-length VEGF165 that is defective in integrin binding by including mutations in the HBD. The full-length VEGF165 mutant defective in integrin binding (R123A/R124A/K125A/K140A/R145A/R149A) was defective in ERK1/2 phosphorylation, integrin β3 phosphorylation, and KDR phosphorylation, although the mutation did not affect KDR binding to VEGF165. We propose a model in which VEGF165 induces KDR (through NTD)-VEGF165 (through HBD)-integrin αvβ3 ternary complex formation on the cell surface and this process is critically involved in potent mitogenicity of VEGF165.

## Introduction

It has been proposed that growth factor signaling requires integrins for cell responses to growth factor ligation of their cognate cell surface receptors [1, 2]. The specific mechanisms and the extent of this integrin-growth factor crosstalk are still not been established. Integrins are cell surface heterodimers that act as receptors for extracellular matrix ligands, cell-surface ligands (ICAM-1, VCAM-1), and soluble ligands that include growth factors [1, 3]. The finding that integrin αvβ3 antagonists inhibit fibroblast growth factor-2 (basic FGF, FGF2) signaling [4] suggested that αvβ3 is involved in growth factor signaling through a crosstalk mechanism [5-7]. We previously reported that FGF1 and FGF2 directly bind to integrin αvβ3, leading to formation of an integrin-FGF-FGFR ternary complex that is required for FGF signaling functions (the ternary complex model) [8-11]. We showed that this model can be applied to other growth factors as well, including IGF1, IGF2, neuregulin-1, fractalkine, and CD40L [12-16]. Thus, direct binding of integrin αvβ3 to growth factors is required for their signaling functions. Notably, we showed that growth factor mutants defective in integrin binding are also defective in signaling functions and can act as antagonists of the signaling process, while the mutants still binding to their cognate receptors [8-11, 13, 14, 16].

Vascular endothelial growth factor (VEGF-A) is a key cytokine in angiogenesis and a major therapeutic target. It has been proposed that VEGF-A signaling requires integrin αvβ3, but it is unclear whether the ternary complex model can be applied, since VEGF-A binding to αvβ3 has not been documented. Among the six main isoforms of VEGF-A, VEGF165 (the 165-amino-acid protein) is the predominant gene product in human tissues and the most potent angiogenesis stimulant [17, 18]. VEGF165 forms homodimers of two anti-parallel monomers that interact via their N-terminal domains which harbor the cognate receptor binding site. VEGF165 exerts mitogenicity by binding to the VEGFR1 (FLT1) and VEGFR2 (KDR) receptor tyrosine kinases [19]. KDR, a 230 kDa glycoprotein, has a lower affinity for VEGF165 (KD=0.75-2 x 10^-10^ M) than VEGFR1 (KD 1-2 x 10^-11^ M). Yet, KDR is the primary mediator of VEGF165 signaling [20]. Within its N-terminal region, KDR has seven IgG-like extracellular domains, of which domains 2 (D2) and 3 (D3) interact with the N-terminal domain of VEGF165 with high affinities [20]. The clinically used anti-angiogenic monoclonal antibody Avastin (Bevacizumab), binds to the N-terminal domain of VEGF165 (residues 1-110) and inhibits VEGF165 binding to KDR. The C-terminal domain of VEGF165 encompasses a heparin-binding domain (HBD, residues 111-165) and a neuropilin-binding site. A plasmin-cleavage site located between the ligand’s N-terminal and heparin binding domains enables proteolytic removal of the HBD, which markedly reduces the mitogenicity of VEGF165 in endothelial cells [21]. This suggests that KDR binding to the N-terminal VEGF165 domain may not be sufficient to exert a cell growth response when the HBD is missing. The reduced mitogenic activity of the VEGF165 homodimer which lacks the HBD is similar to that observed for VEGF121, a splice variant that lacks exons 6a and 7 which encode most of the HBD. Although evidence suggests that the HBD plays a role in VEGF165 dependent processes including angiogenesis, it is unclear how the HBD contributes to VEGF165 mediated cell responses.

In the present study we first showed that αvβ3 directly binds to HBD. We then identified amino acid residues critical for αvβ3 by docking simulation of interaction between HBD and αvβ3, and subsequent mutagenesis studies. Our results indicate that integrin αvβ3 binds to the HBD of VEGF165 and leads to formation of a ternary complex between αvβ3, VEGF165 and KDR. Having identified the integrin binding site within the VEGF165-HBD, our studies using VEGF165-HBD mutants defective in integrin binding suggest that αvβ3 binding to VEGF165 is essential for KDR-mediated VEGF165 cell activation. The results from our work will enable development of new synergistic therapies that co-target αvβ3-and KDR-VEGF interaction to control diseases that involve aberrant cell proliferation and angiogenesis.

## Materials and Methods

Recombinant soluble αvβ3 was synthesized in Chinese hamster ovary (CHO) K1 cells using the soluble αv and β3 expression constructs and purified by nickel-nitrilotriacetic acid (Ni-NTA) affinity chromatography as described [22].

### Plasmid Construction and protein purification

The cDNA fragment of the N-terminal domain (NTD) and the C-terminal domain (CDT) of VEGF165 were amplified using primers with human VEGF165 cDNA (Open Biosystems, Lafayette, CO) as a template, and was subcloned into the BamHI/EcoRI site of the PET28a expression vector. The cDNA fragment of domain 1 (D1) of KDR was amplified using full-length human KDR cDNA as a template and subcloned to the BamH1/EcoR1 site of pET28a. The insoluble protein was synthesized in BL21 induced by IPTG and was solubilized in 8 M urea, purified by Ni-NTA affinity chromatography and refolded as described [12]. KDR D1 cDNA was also subcloned to the BamH1/EcoR1 site of pGEX2T vector and protein was synthesized in BL21 induced by IPTG and KDR D1 fused to GST was purified from cell lysates by glutathione-affinity chromatography.

### ELISA-type Analysis

Ninety-six-well plates were coated with KDR D1 or D1D2 in a 10 μg/mL concentration and incubated at 37°C for one hour. The wells were further with 0.1% BSA/PBS boiled at 80°C for 20 minutes then cooled before application of 300 μl/well. After 1 hour of blocking, protein was serially diluted in PBS, 50 μl/well, for 1 hour at room temperature. After washing with 0.05% Tween 20/PBS, an anti-His was applied for one hour at room temperature. Three times of washes were performed with 0.05% Tween20/PBS before detection with 3,3’,5,5’-Tetramethylbenzidine (TMB) solution, 100 μl/well. The reaction was stopped by application of 2N HCl, 50 μl/well, and measurements were made at OD=450 nm with a plate reader.

### Surface plasmon resonance study

KDR D1 (His-tagged) was immobilized on the CM5 sensor chip using a standard amine coupling procedure in HBS-EP buffer (0.01 M Hepes, pH 7.4, 0.15 M NaCl, 3 mM EDTA, and 0.0005% of surfactant P20), and the same buffer. Samples were injected at 50 μl/min for 1.8 min. The HBS-EP buffer was then injected at 50 μl/min for 3 min to allow the bound VEGFs to dissociate from the KDR D1.

### Binding of soluble αvβ3

A 96-well plate was coated with PBS-diluted protein and incubated at 37°C for one hour. The wells were further blocked with 0.1%BSA/PBS boiled at 80°C for 20 min then cooled down before application of 300 μl/well. After half an hour of incubation at room temperature, 1×10^5^ cells/well in DMEM or Hepes-Tyrode buffer+ 1mM MgCl_2_/1mM MnCl_2_ were added and the plate was further incubated in a 5% CO_2_ containing incubator at 37°C for one hour. Cells were removed and washed with dilution buffer A Phosphatase buffer (100μl/well) was applied, and the plate was incubated at 37°C for 45 min. 1 M NaOH (50μl/well) was applied to stop the reaction and the plate was read at OD=405 nm by the plate reader.

### Docking simulation

Docking simulation of interaction between the HBD of VEGF165 (2VGH.pdb), which does not contain N-terminal KGD motif, and integrin αvβ3 was performed using AutoDock3 as described [12]. In the present study, we used the headpiece (residues 1-438 of αv and residues 55-432 of β3) of αvβ3 (open-headpiece form, 1L5G.pdb). Cations were not present in αvβ3 during docking simulation [8, 23].

### SDS-PAGE analysis of wild type and mutant VEGF165

In mutant VEGF165 all six amino acids within the HBD identified as required for integrin αvβ3 binding were changed to Ala (R123A/R124A/K125A/K140A/R145A/R149A). Molecular size values in kDa.

### Pull-down of VEGF165 by KDR

His-tagged KDR was immobilized on Ni-NTR beads and incubated with full-length VEGF165 wild type (WT) vs VEGF165 mutant (mut) protein in binding buffer for 2 h at 4C, before elution and analysis of the retained protein by SDS-PAGE. Both wild type and mutant VEGF165 bind to KDR.

### ERK1/2 activation in HUVECs by VEGF165

Starved HUVECs in M200 basal medium without low serum growth supplement were treated with VEGF165 (10 ng/ml) for 10 min before lysis and Western blot analysis. Wild type (WT) but not mutant (mut) VEGF165 activates ERK1/2 phosphorylation in human umbilical vein endothelial cells (HUVEC).

### Integrin αvβ3 phosphorylation in HUVEC in response to VEGF165

Starved HUVECs were treated with VEGF165 wild type vs mutant (10 ng/ml) before lysis and Western blot analysis of β3 integrin subunit phosphorylation. Wild type (WT) but not mutant (mut) VEGF165 activates integrin αvβ3 phosphorylation in HUVECs.

### KDR Y1175 phosphorylation in HUVEC in response to VEGF165

In HUVECs, wild type (WT) but not mutant (mut) VEGF165 activates KDR Y1175 phosphorylation known to stimulate endothelial cell proliferation and migration. Starved HUVECs were treated for Western blot analysis.

### Statistical Analysis

We reported results as the mean ± standard error of the mean and performed calculations using Prism 7. Statistical analysis was performed using ANOVA.

## Results

### Direct binding of integrin αvβ3 to the VEGF165 heparin binding domain (HBD)

It has been proposed that VEGF165 interacts with αvβ3, but VEGF121 which lacks the HBD, does not [24]. This implies that the HBD likely contains a binding site for αvβ3, however, the specifics of an HBD-αvβ3 interaction are unclear. We found that soluble αvβ3 bound to immobilized VEGF165-HBD protein in a dose-dependent manner, but not to the N-terminal domain (NTD) in an ELISA-type binding assay (Fig. 1a,b). This binding was suppressed by heat treatment of the HBD, suggesting that the interaction requires proper folding of the HBD. This binding was inhibited by function blocking mAb 7E3 specific to β3 integrin (Fig. 1c), which is mapped in the classical ligand-binding site within the β3 integrin subunit [25, 26]. Binding of soluble αvβ3 to the HBD is cation dependent (Mn^2+^>Mg^2+^=Ca^2+^>EDTA) (Fig. 1d). In a surface plasmon resonance study, the K_D_ of HBD-αvβ3 interaction was calculated as 4.7 x 10^-7^ M (Fig. 1e).

**Fig. 1.**
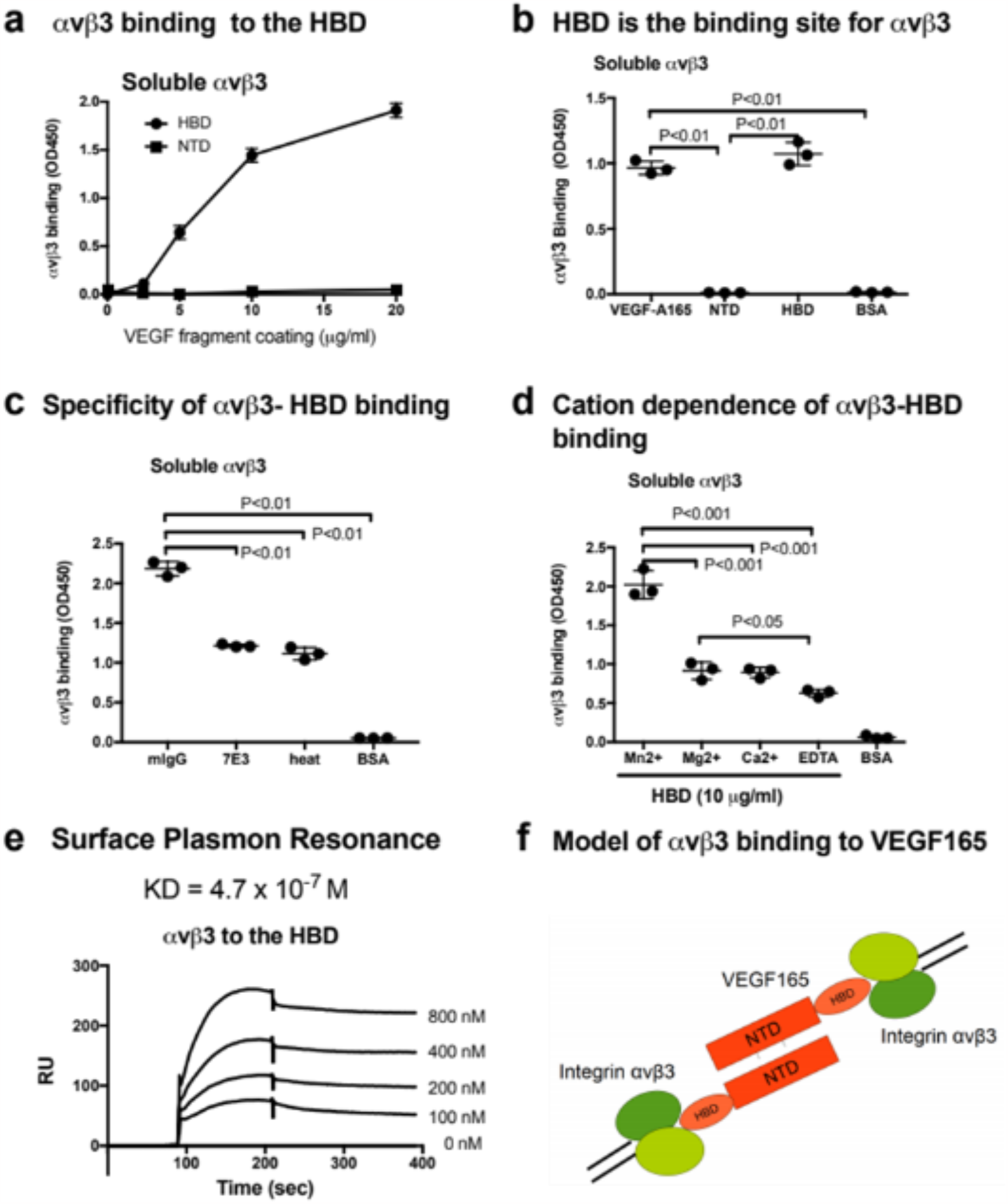
Integrin αvβ3 binds to the heparin binding domain (HBD, residues 111-165) of VEGF165, but not to the N-terminal domain (NTD, residues 1-110). a, b: Interaction of soluble αvβ3 with immobilized VEGF165 HBD vs NTD. Microtiter wells were coated with increasing concentrations of VEGF165 or its fragments and incubated with soluble αvβ3, after blocking non-specific protein binding sites with BSA. In (b) control wells were coated with BSA. αvβ3 binding was detected with non-function blocking anti-β3 mAb AV10. c: Specificity of αvβ3 binding to the VEGF165-HBD documented based on inhibition of αvβ3 binding by function-blocking anti-β3 antibody, mAb 7E3, or heat treatment, but not by control mouse IgG (mIgG). Control wells were coated with BSA. d: Cation-dependence of αvβ3 binding to the VEGF165-HBD. Binding of soluble αvβ3 to HBD protein (10 μg/ml) immobilized on microtiter plates and blocked with BSA, in the absence of divalent cations (EDTA) or presence of Mn^2+^, Ca^2+^, or Mg^2+^ (1 mM). Control wells were coated with BSA. e: Surface Plasmon Resonance analysis of αvβ3-HBD interaction. αvβ3 protein was immobilized to a sensor chip and binding of solubilized mobile HBD protein was measured as analyte at increasing concentrations. f: Model of αvβ3 binding to the C-terminal HBD within a VEGF165 homodimer formed by two anti-parallel VEGF164 monomers known to interact via their N-terminal domains (NTD). The NTD does not contribute to αvβ3 binding as documented in our study. Where applicable, data are shown as means +/-SD of triplicate experiments.

These results suggest that the VEGF165 HBD is a specific ligand for αvβ3 (Fig. 1f). These findings predict that the VEGF165 binds to αvβ3 through the HBD and to KDR via the N-terminal domains (NTD), resulting in the integrin-VEGF165-KDR ternary complex.

### Mapping amino acid residues within the VEGF165 HBD for integrin binding

To predict which amino acid residues are involved in αvβ3 binding, we performed docking simulations between the HBD (2VGH.pdb) and αvβ3 (1L5G.pdb, with open headpiece) using Autodock3 (Fig. 2a). The simulation predicted that the HBD binds to αvβ3 at high affinity (docking energy at −24.6 Kcal/mol). Amino acid residues predicted as involved in the HBD-αvβ3 interaction are Arg-123, Arg-124, Lys-125, Lys-140, Arg-145, and Arg-149. We selected several Arg or Lys residues of the HBD in the predicted binding interface for mutagenesis (Fig. 2b). Binding of soluble αvβ3 to immobilized HBD wild type vs mutant protein was measured by ELISA-type binding assays. The Arg123/Arg124/Lys125 to Ala (R123A/R124A/K125A) mutation and the Lys140/Arg145/Arg149 to Ala mutation (K140A/R145A/R149A) partially suppressed the binding of αvβ3 to wild type HBD (Fig. 2c). The combined R123A/R124A/K125A/K140A/R145A/R149A mutation nearly completely blocked integrin binding (Fig. 2d). These results indicate that these amino acid residues are critical for VEGF165 HBD binding to αvβ3.

**Fig. 2.**
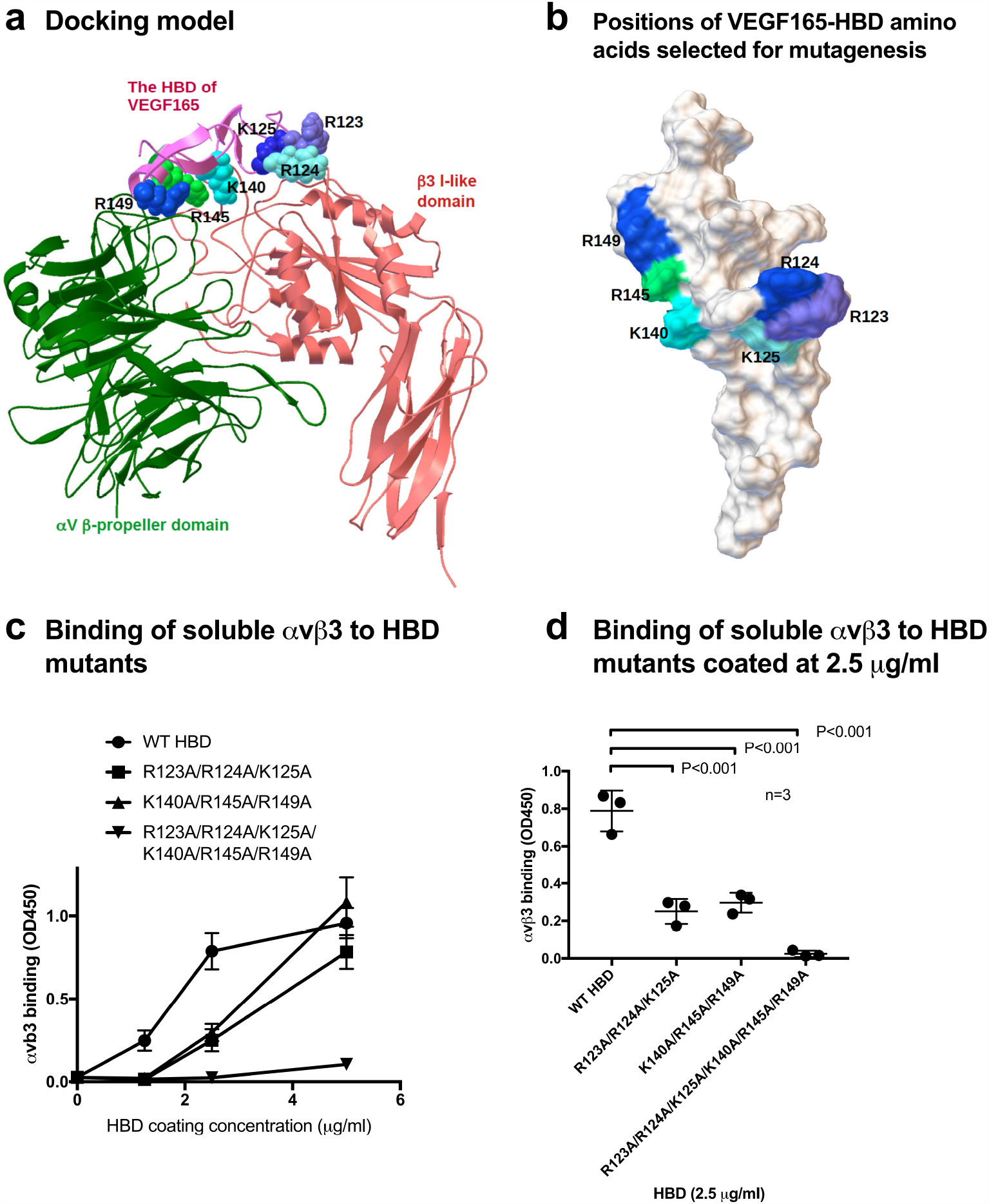
Mapping the integrin binding sites within the heparin-binding domain (HBD) of VEGF165. a: Docking model of the interaction between integrin αvβ3 and the HBD based on docking simulation using Autodock 3. HBD amino acid residues predicted to contribute to αvβ3 binding are Arg-123, Arg-124, Lys-125, Lys-140, Arg-145, and Arg-149. b: Position of HBD amino acids within the predicted integrin binding interface were selected for mutagenesis and changed to Ala in combinations. c: Binding of soluble αvβ3 to VEGF165-HBD mutants coated onto microtiter wells at increasing concentrations. Non-specific binding sites were blocked with BSA. d: Binding of soluble αvβ3 to VEGF165-HBD mutants, coated at near saturation concentration (2.5 μg/ml) identified for the wild type (WT) HBD protein, revealed that all HBD amino acids predicted to contribute to αvβ3 binding are required for VEGF165-integrin αvβ3 interaction. Where applicable, data are shown as means +/-SD of triplicate experiments.

### Full-length VEGF165 with a mutated HBD defective in αvβ3 binding is defective in signaling functions while still binding KDR

Full-length VEGF165 protein with the combined HBD R123A/R124A/K125A/ K140A/R145A/R149A mutations (referred to as VEGF165-HBD mutant) (Fig. 3a) was synthesized in E. coli as insoluble protein, purified under denaturation, refolded and further purified by FPLC gel-filtration. The protein migrated as a single band with expected size (Fig. 3a). The protein bound to immobilized KDR in pull-down assays (Fig. 3b), indicating that KDR binding specificity was retained. The VEGF165-HBD mutant lacked integrin binding αvβ3 -as expected. Used as a ligand in human endothelial cell (HUVEC) cultures, VEGF165-HBD mutant protein failed to induce ERK1/2 activation and integrin β3 phosphorylation (Fig. 3c-e). Furthermore, VEGF165-HBD mutant protein also failed to activate KDR Y-1175 phosphorylation in HUVECs (Fig. 3f), despite the ability of the mutant protein to bind KDR (Fig.3b). These findings indicate that binding of integrin αvβ3 to the HBD of VEGF165 is critical for signaling functions of VEGF165.

**Fig. 3.**
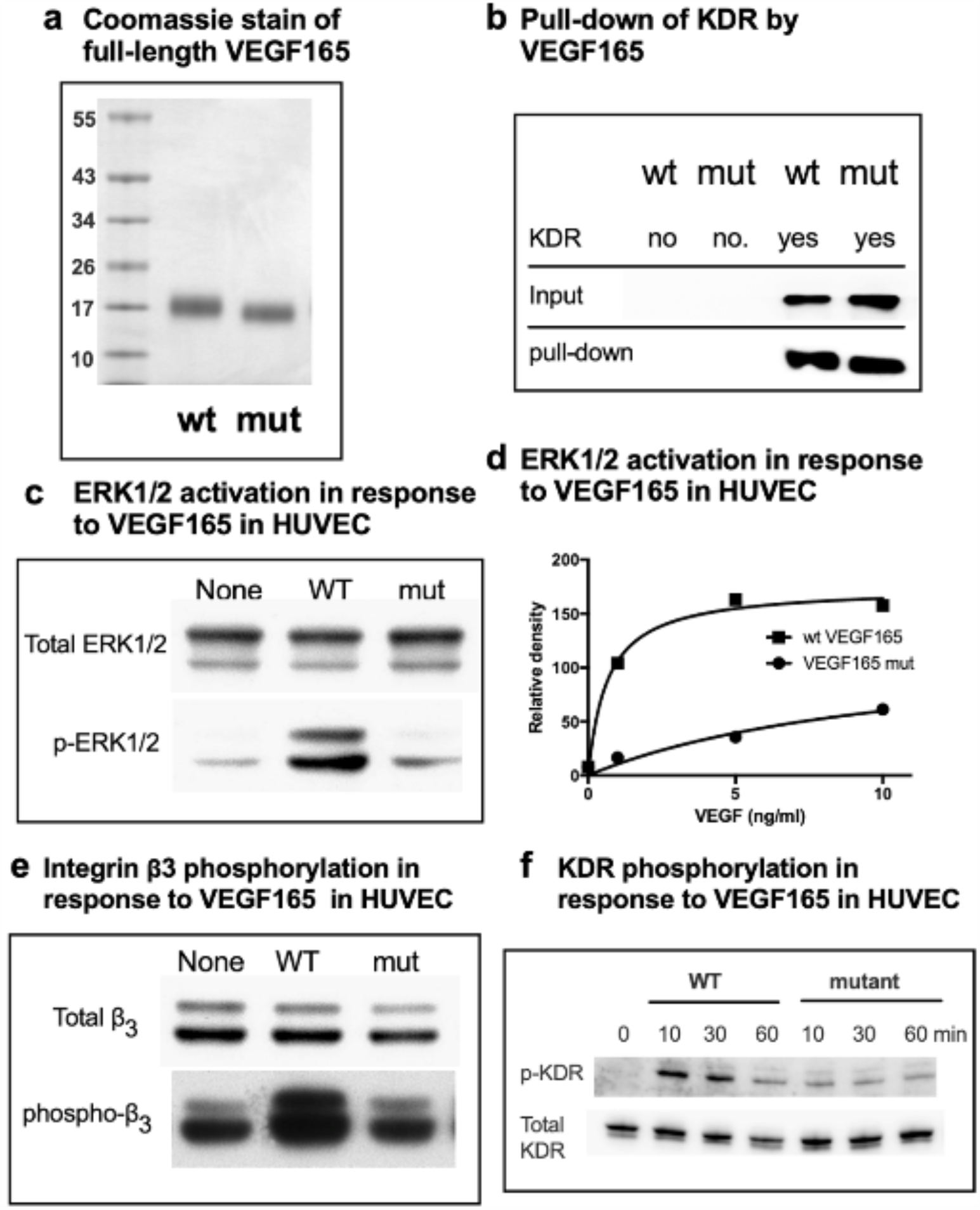
Partial characterization of the full-length VEGF165 mutant defective in integrin binding. a: SDS-PAGE analysis of wild type and mutant VEGF165. In mutant VEGF165 all six amino acids within the HBD identified as required for integrin αvβ3 binding were changed to Ala (R123A/R124A/K125A/K140A/R145A/R149A). Molecular size values in kDa. b: Pull-down of VEGF165 by KDR. His-tagged KDR was immobilized on Ni-NTR beads and incubated with full-length VEGF165 wild type (WT) vs VEGF165 mutant (mut) protein in binding buffer for 2 h at 4C, before elution and analysis of the retained protein by SDS-PAGE. Both wild type and mutant VEGF165 bind to KDR. c: ERK1/2 activation in HUVECs by VEGF165. Starved HUVECs in M200 basal medium without low serum growth supplement were treated with VEGF165 (10 ng/ml) for 10 min before lysis and Western blot analysis. Wild type (WT) but not mutant (mut) VEGF165 activates ERK1/2 phosphorylation in human umbilical vein endothelial cells (HUVEC). d: Dose dependence of ERK1/2 activation in HUVECs by VEGF165. e: Integrin αvβ3 phosphorylation in HUVEC in response to VEGF165. Starved HUVECs were treated with VEGF165 wild type vs mutant (10 ng/ml) before lysis and Western blot analysis of β3 integrin subunit phosphorylation. Wild type (WT) but not mutant (mut) VEGF165 activates integrin αvβ3 phosphorylation in HUVECs. f: KDR Y1175 phosphorylation in HUVEC in response to VEGF165. In HUVECs, wild type (WT) but not mutant (mut) VEGF165 activates KDR Y1175 phosphorylation known to stimulate endothelial cell proliferation and migration. Starved HUVECs were treated for Western blot analysis as in panel e.

Our previous studies showed that mutants of growth factors (e.g., FGF1, IGF1, and fractalkine) that lack integrin binding, still bound to their cognate receptors and can act as dominant-negative antagonists [8-11, 13, 14, 16]. We therefore hypothesize that the VEGF165-HBD mutant defective in integrin binding can act as an antagonist for KDR-VEGF165 mediated cell responses and reveal target information for novel synergistic therapeutic approaches.

## Discussion

The present study establishes that integrin αvβ3 binds to the isolated HBD, not NTD, of VEGF165, indicating that the HBD is a specific ligand for αvβ3. This is consistent with a previous report that the HBD contains an αvβ3-binding site [24]. The proteolytic removal of the HBD markedly reduced the mitogenicity of VEGF165, suggesting that the binding of the NTD to KDR D2D3 may not be sufficient for potent mitogenicity [21, 27]. The present results suggest that VEGF165 bind to integrin αvβ3 via the HBD and to KDR via NTD and as a result induces integrin-VEGF165-KDR ternary complex formation on the cell surface.

The present study identified amino acid residues in the HBD that are critical for integrin binding (Arg-123, Arg-124, Lys-125, Lys-140, Arg-145, and Arg-149) using docking simulation and mutagenesis. Mutations R123A/R124A/K125A and K140A/R145A/R149A effectively reduced integrin binding. Combined mutations of all these amino acids (R123A/R124A/K125A/K140A/R145A/R149A) almost completely suppressed the binding of αvβ3 to the HBD. We generated full-length VEGF165 with the combined HBD mutations (R123A/R124A/K125A/K140A/R145A/R149A) did not affect KDR binding, but effectively suppressed ERK1/2 activation, phosphorylation of integrin β3, and KDR phosphorylation in HUVEC. These findings suggest that the binding of VEGF165 to αvβ3 through the HBD plays a critical role in VEGF165 signaling. We propose that the ternary complex model can be applied to VEGF165.

We have reported that FGF1 and FGF2 directly bind to integrins and these interactions lead to ternary complex formation (integrin-FGF-FGFR), which is required for their signaling functions (the ternary complex model) [8-11]. This suggests that direct binding of integrins to FGF that positively regulates FGF signaling is required. Consistently, mutants of FGF defective in integrin binding are functionally defective and suppress signaling induced by WT FGF (dominant-negative antagonists) [9, 10]. These findings are consistent with the reports that antagonists to integrins (e.g., αvβ3) block angiogenesis induced by FGF2 [4]. We have also reported that integrins also directly bind to several growth factors other than FGF (IGF-1 and −2, neuregulin-1, fractalkine, and CD40L) and positively regulate their signaling functions [12-16, 28]. We showed that growth factor mutants defective in integrin binding (e.g., IGF1, IGF2, fractalkine and CD40L) acted as dominant-negative antagonists [13, 14, 28]. We thus predict that VEGF165 mutant (R123A/R124A/K125A/K140A/R145A/R149A) act as a dominant-negative antagonist.

## Acknowledgements

This project was supported by pilot funding from the Comprehensive Cancer Center at UC Davis School of Medicine. This work is partly supported by the UC Davis Comprehensive Cancer Center Support Grant (CCSG) awarded by the National Cancer Institute (NCI P30CA093373) and the Pilot funding from the Department of Dermatology.

## Author Contributions

YK Takada, J. Yu, X. Ye, and C-Y Wu: data curation.

M.Fujita., and B.H. Felding, resources, writing—review and editing

Y Takada: conceptualization, formal analysis, funding acquisition, and project administration.

## Conflict of Interest Statement

The authors declare that they have no conflict of interest.

**Table 1.**
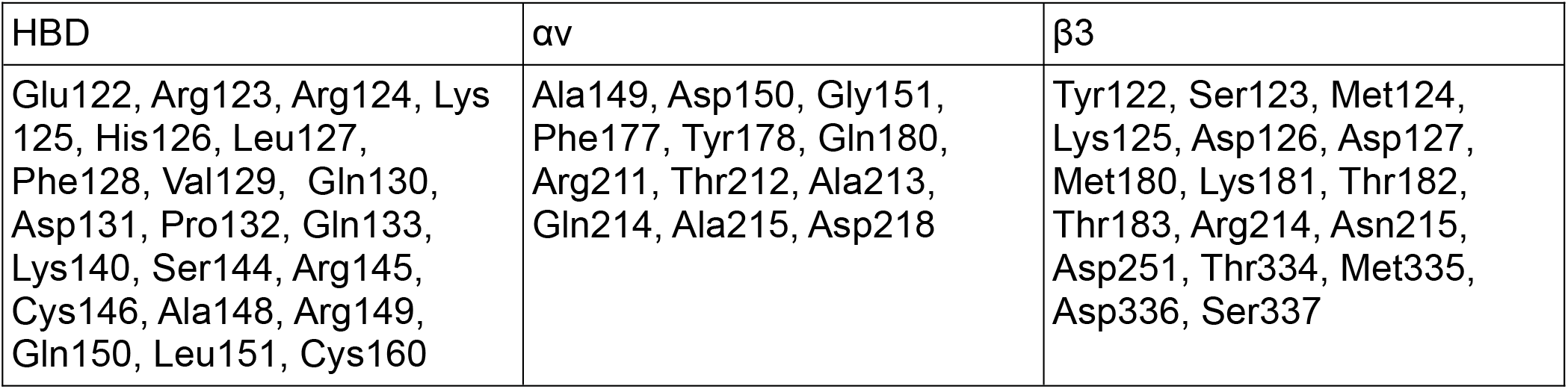
Amino acid residues predicted to be involved in HBD binding to integrin αvβ3. Amino acid residues within 0.6 nm between the HBD and αvβ3 were selected using Swiss PDB Viewer (version 4.1) (Swiss Institute of Bioinformatics, Basel, Swiss).

| HBD | $\alpha\text{v}$ | $\beta 3$ |
| --- | --- | --- |
| Glu122, Arg123, Arg124, Lys125, His126, Leu127, Phe128, Val129, Gln130, Asp131, Pro132, Gln133, Lys140, Ser144, Arg145, Cys146, Ala148, Arg149, Gln150, Leu151, Cys160 | Ala149, Asp150, Gly151, Phe177, Tyr178, Gln180, Arg211, Thr212, Ala213, Gln214, Ala215, Asp218 | Tyr122, Ser123, Met124, Lys125, Asp126, Asp127, Met180, Lys181, Thr182, Thr183, Arg214, Asn215, Asp251, Thr334, Met335, Asp336, Ser337 |

